# Variable fluid mechanics explain why static efficacy tests overestimate sanitizer performance against Listeria

**DOI:** 10.64898/2026.05.13.724842

**Authors:** Yang Jiao, Jakob Baker, Calvin Slaughter, Devin Daeschel, Abigail B. Snyder

## Abstract

Pathogen cross-contamination during food production is primarily controlled through environmental sanitation. However, sanitizer efficacy is often studied in bench-scale experiments that poorly approximate the fluid dynamics of sanitization and limits our understanding of commercial sanitization efficacy. This study paired computational fluid dynamics (CFD) estimates of shear stress with experimental measurements of *Listeria innocua* reduction on stainless steel following treatment with 100 ppm hypochlorite sanitizer. At the pilot-scale, sanitizer spray manually applied by researchers achieved a 2.6 ± 0.4 log CFU/surface reduction; however, microbial reduction from manual operation of sanitizer spray equipment differed significantly between researchers (p < 0.05). Microbial reduction varied by location following stationary, bench-scale spray application of sanitizer for 3 s. The greatest reduction was at the point of sanitizer spray impingement (7.5 ± 0.5 log CFU/surface) and directly adjacent to the impingement point (6.4 ± 0.7 log CFU/surface) where shear stress was the highest. Significantly less microbial reduction (0.4 ± 0.1 log CFU/surface) occurred where shear stress was lowest in the fluid-film of sanitizer running down from the impingement point (p < 0.05). Static submersion of inoculated coupons in sanitizer for 3 s resulted in a log reduction of 2.3 ± 0.1 log CFU/surface. Discrepancies between bench-scale spraying, pilot-scale spraying, and submerged coupons demonstrate the need for sanitizer efficacy testing under realistic conditions to better estimate the risk reduction achieved through sanitation programs.

**IMPORTANCE:** Sanitation is critical for controlling pathogen cross-contamination during food production. These findings highlight the limitations of traditional approaches to sanitizer efficacy testing, not because they are invalid, but because they do not reflect the level of microbial reduction typically achieved in application. We demonstrate that these differences in outcomes are attributable to fluid dynamics and exposure, which are not well approximated in submerged coupon experiments. Accurate estimation of microbial reduction from sanitizer application is needed to guide food safety policy decisions. For example, overestimation of the risk reduction conferred by sanitizer treatment may result in food safety policies that neglect other sources of microbial reduction within sanitation programs.

## 1. INTRODUCTION

Cross-contamination of pathogens from environmental surfaces to food during processing has contributed to numerous recalls and foodborne illness outbreaks^1–3^. Pathogens, including *Listeria monocytogenes* can persist on environmental surfaces despite sanitation which is the primary control against cross-contamination of food from environmental pathogens^1,4–7^. Environmental surfaces in food manufacturing plants include the food-contact and non-food-contact surfaces of processing equipment and utensils, non-processing equipment and tools such as push carts and floor mats, as well as the building’s structures such as floors and drains^8^. Apart from the internal surfaces of equipment that can be cleaned using automated clean-in-place (CIP) systems, most food-contact and environmental surfaces are cleaned manually through wiping, spraying, or washing, and are subsequently treated with an aqueous sanitizer spray.

It is commonly stated that the purpose of cleaning is to remove food residues, while the purpose of sanitizing is to chemically inactivate any remaining microbial cells on surfaces^3^. However, this framework oversimplifies sanitation by overstating the role of chemical inactivation and neglecting the fact that a significant degree of microbial reduction is due to physical removal via fluid dynamic forces such as shear stress^9^. This conceptual simplification has carried over into sanitizer efficacy testing, which largely evaluates chemical lethality under static conditions and does not account for microbial reduction through physical removal during spray application.

The efficacy of surface sanitization through sanitizer spraying is dependent on physical treatment variables such as the application time (the amount of time the sanitizer treatment is actively applied), contact time (the amount of time the sanitizer is allowed to remain on surfaces before washing), shear stress (the frictional force of liquid sanitizer moving over a surface), and mode of application (e.g. wiping, spraying, flooding)^9–11^. In practice, sanitizing surfaces is a dynamic process with variable shear stress and contact time. Some surface locations receive the full force from the sanitizer spray directly at the point of surface impingement. Locations adjacent to the impingement point are exposed to sanitizer that splashes or flows outward with diminished force. Other locations are exposed to sanitizer that flows down from the point of impingement in a fluid-film due to gravity, providing even lower shear forces on the surface. Furthermore, since sanitizer sprayers are manually operated by personnel, the application of the treatment varies depending on individual techniques, including differences in spray duration, nozzle proximity to surface, and consistency at each location on the surface.

Despite this inherent variability in physical exposure conditions during spray-based sanitation, most sanitizer efficacy testing frameworks do not explicitly account for the influence of shear stress or fluid dynamics on microbial reduction. This is reflected in the Environmental Protection Agency’s methodology for validating sanitizers for food-contact surface use, which requires sanitizers to achieve a ≥5-log reduction of Escherichia coli (ATCC 11229) and Staphylococcus aureus (ATCC 6538) suspended in static sanitizer solutions within ≤30 s^12^. Similar to the EPA’s sanitizer testing methodologies, prior bench-scale studies evaluating sanitizer efficacy have frequently neglected the role of shear stress in microbial reduction, instead relying on immersion of inoculated surfaces in sanitizer solutions^13,14^. Likewise, pilot-scale challenge studies that manually apply sanitizer sprays to inoculated surfaces have not quantified how microbial reduction varies with the range of shear stress exposures at different locations across a surface^15,16^. Collectively, these approaches to measuring sanitizer efficacy de-emphasize the role of the spatially variable fluid dynamics associated with sanitizer sprays, thereby limiting interpretation of how physical removal contributes to total microbial reduction.

A more representative evaluation of sanitizer efficacy requires explicitly accounting for shear stress and fluid dynamic conditions alongside chemical action. By integrating these physical factors into sanitizer efficacy assessment, it becomes possible to more accurately characterize microbial reduction under realistic sanitation conditions and to generate data that better inform sanitation practices and risk mitigation strategies in food manufacturing facilities.

## 2. MATERIALS AND METHODS

### 2.1 Inoculum preparation

Three strains (ATCC 51742, FSL C2-008, FSL A5-0424) of *Listeria innocua* were selected for this study from the Food Safety Laboratory at Cornell University (Ithaca, NY) based on prior studies with surface detachment^17–19^. Each strain was stored at -80°C in 15% glycerol (v/v). A loopful (10 µL) of frozen stock for each strain was cultured in brain heart infusion (BHI) broth (BD, Thermo Fisher Scientific, Waltham, MA, USA) by incubation at 37±2°C for 24±3 h with three successive transfers (24 h intervals), then streaked onto BHI agar plates and incubated at 37±2°C for 24 h. An isolated colony from each plate was inoculated into 2 × 5 mL BHI broth and incubated at 37 ± 2°C for 24 h. The culture was centrifuged at 5,000 RPM for 10 min (Eppendorf 5804R, Eppendorf, NY, USA), and the pellet was washed twice with 0.1% phosphate buffered saline (PBS) (BD, Thermo Fisher Scientific, Waltham, MA, USA). After washing, the cell pellets were resuspended in 0.5 mL of 0.1% phosphate buffered saline to achieve a cell concentration of ∼8-log CFU/mL. A 3-strain cocktail was prepared by mixing equal volumes of the three cultures. A cocktail was used instead of individual strains to provide a conservative assessment, as combining multiple *Listeria* strains results in the most conservative strain dominating the observed response.

### 2.2 Sanitizer preparation

Sanitizer solutions containing 100 ppm available chlorine were prepared by diluting tap water with XY-12 (Ecolab, St. Paul, CA, USA), a chlorine-based liquid sanitizer containing 8.4% sodium hypochlorite. If present, resistant organisms in the tap water would be expected at low levels unlikely to influence study outcomes. Consistent with this, Mazoua and Chauveheid (2005) reported that most post-treatment water samples contained fewer than 10 CFU/L of aerobic spore-forming bacteria^20^. The sanitizer concentration was verified using a chlorine test kit (FryOilSaver Co., Matthews, NC, USA). The use of this product and a target concentration of 100 ppm sodium hypochlorite was selected based on its application in food processing environments^21^.

### 2.3 Surface preparation

For the pilot-scale study, a stainless steel table with a top surface with a *length* × *width* of 120 × 70 cm^2^ was set on its side perpendicular to the floor. The stainless steel table was uniformly sprayed with ethanol (70% EtOH) and wiped with paper towels, then air-dried for 10 min. For the bench-scale spray and submersion experiments stainless steel sheets (0.46 mm, 316/316 L, cold roll 2B) were cut into coupons with dimensions of 30 × 30 cm^2^, and 10 × 10 cm^2^ respectively by the Machine Shop at Cornell University. Different coupon sizes were selected to allow for complete submersion of the coupon for the submersion trials and for site specific inoculation and recovery in the bench-scale spray chamber experiments. All coupons were sterilized in an autoclave (Steris®, Mentor, OH, USA) at 121°C for 30 min under 15 psi. Stainless steel surfaces were selected because they are common food contact materials in manufacturing facilities and are routinely subjected to sanitizer spray applications.

### 2.4 Pilot-scale sanitation experiment

#### 2.4.1 Pilot-scale surface inoculation

The *L. innocua* cocktail was inoculated (10 μL spot × 10 spots) on the top of the stainless steel tables while they were in an upright position, yielding approximately 7 log CFU/surface. The spot locations were predetermined and consistent among replicates. Inoculated surfaces were allowed to air dry at 22°C for 3 h while upright, then placed perpendicular to the floor for treatment. All treatments were conducted in biological triplicate.

#### 2.4.2 Treatment of surfaces by manual spray application

Three researchers were familiar with industrial sanitation and were instructed to spray sanitizer (100 ppm sodium hypochlorite) on a test surface using a pressurized sprayer (FI-5N, Foamit, Grand Rapids, MI, USA) with a flat-fan shape VeeJet nozzle (H1/4U-SS2520). The sprayer was connected to a ∼19 L (5 gallon) reservoir with a 5 m hose as well as a compressed air system with 50 psi pressure. They were told to assume that they were sanitation personnel in a food processing environment. No prior knowledge of the inoculation sites was given. The researchers were informed to tie back all hair, wear long pants, closed-toed shoes (not cloth shoes), as well as don splash-proof lab goggles, nitrile gloves, and a lab coat. No additional instructions were provided besides the safety protocol for manual operation of the sprayer. The distance (cm) between the sprayer nozzle and surface, the duration (s) of spraying treatment, and the application path of the sanitizer spray on the inoculated surface were extracted from video recordings of the treatments using an Apple iPhone XR (Apple Inc., Cupertino, CA, USA). Each researcher determined their own spray path, treatment duration, and nozzle distance from the surface; however, these variables were approximately the same for each researcher among their three replicate treatments. A 120 s sanitizer contact time was established between the end of each researcher’s spray treatment and before recovery and enumeration.

### 2.5 Bench-scale sanitizer spray-chamber experiment

#### 2.5.1 Bench-scale spray-chamber construction

The sprayer, nozzle, hose, reservoir and air compressor used for manual spray application were used for bench-scale spray application as well. A bench-scale sanitation chamber (38 × 38 × 38 cm^3^) was custom built for sanitizer spray application onto a vertically fixed 30 × 30 cm^2^ surface. The frame was made with Polyvinyl Chloride (PVC) tubes and wrapped with PVC plastic sheets to enclose the spray from splashing to the surrounding environment. The enclosure had an opening at the front and a PVC tube was attached horizontally approximately 24 cm from the frame’s bottom to secure the sprayer’s position, ensuring precise control and consistent replicates during the spraying experiments. A manufactured PVC coupon (30 × 30 cm^2^) was tied at the back of the frame to secure interchangeable stainless steel coupons (30 × 30 cm^2^) at a distance of 38 cm directly in front of the sprayer nozzle.

#### 2.6.2 Inoculation of surfaces for bench-scale spray-chamber experiments

Three locations on a stainless steel coupon (30 × 30 cm^2^) were selected as inoculation sites for the bench-scale sanitizer spraying experiment. The locations were named as “Impingement,” “Impingement-adjacent,” and “Fluid-film,” indicating their position relative to the sanitizer spray’s impingement point on the surface. During spraying, the “Impingement” sampling location was at the center (0 cm) of the spray impingement on the surface, the “Impingement-adjacent” sampling position was 5.1 cm (2 in) to the left of the “Impingement” sampling location. The “Fluid-film” location was 7.6 cm (3 in) below the “Impingement” sampling location. The surface was spot inoculated (0.1 mL; 7 log CFU) at each location and then the inoculated surface was placed in a biosafety hood (Thermo Fisher Scientific, Waltham, MA, USA) (22°C) to dry down for 3 h prior to treatment. All treatments were conducted in biological triplicate.

#### 2.5.3 Sanitation of surfaces in a bench-scale spray-chamber

A bench-scale study was conducted by applying the prepared sanitizer using a fixed-position sprayer oriented perpendicular to vertically mounted surfaces. The sprayer was operated for 3 s at a distance of 38 cm from the surface. This application time and distance were selected to approximate a brief, targeted spray commonly used during routine sanitation or corrective cleaning events in food processing environments. Additional experiments were conducted to evaluate the influence of application duration and contact time on sanitizer efficacy. Specifically, treatments included: (1) extending the spray application from 3 s to 2 min to represent the impact of spray-based sanitation, and (2) maintaining a 3 s spray application followed by a contact time of 2 min prior to neutralization in accordance with sanitizer application instructions to simulate situations in which surfaces remain wet with sanitizer for an extended period following its application.

### 2.6 Sanitizer submersion of coupons experiment

#### 2.6.1 Inoculation of coupons for submersion

Cocktails were spot inoculated (0.1 mL; 7 log CFU) onto the center of stainless steel coupons (10 × 10 cm^2^). Then, the inoculated coupon was placed in a biosafety hood (Thermo Fisher Scientific, Waltham, MA, USA) (22°C) to dry down for 3 h prior to treatment. All experiments were conducted in biological triplicate.

#### 2.6.2 Treatment of coupons by submersion into static sanitizer

Coupons were submerged for 3 s or 120 s of application time in 1 L of prepared 100 ppm sodium hypochlorite sanitizer. Following treatments that did not include a subsequent hold period for extended contact time after removal from the sanitizer, coupons were immediately processed according to the steps in *2.7 Recovery and enumeration*. Following treatments with subsequent hold periods to extend contact times, coupons were stored vertically for 120 s before being processed according to the steps in *2.7 Recovery and enumeration*.

### 2.7 Recovery and enumeration

Surfaces were swabbed with a sponge-stick with 10 mL Dey/Engley (D/E) neutralizing broth (SSL10DE, Neogen, St Paul, USA). Swabbing was conducted both vertically and horizontally (5 top to bottom strokes, followed by 5 left to right strokes) as recommended by the swab’s manufacturers (Neogen, St. Paul, USA). For the bench scale and pilot scale trials, swabbing was conducted in 10 x 10 cm^2^ square centered on the point of inoculation. At the coupon scale, the entire surface of the coupon was swabbed. This swabbing protocol was validated in preliminary studies to ensure consistently accurate microbial counts. The swabs were stomached for 2 min at 260 rpm (Stomacher® 400 Lab Blender Series, Seward, UK). Tenfold serial dilutions were conducted, then 100 μL of dilutions were spread onto BHI plates and incubated at 37°C for 24 h before being enumerated. Recovered colonies were presumptively attributed to L. innocua because surfaces were decontaminated prior to inoculation, and preliminary post-disinfection surface swabs yielded counts below the limit of detection on BHI.

### 2.8 Computational fluid dynamics modeling

A computational fluid dynamics model (CFD) was constructed in ANSYS Fluent (ANSYS, 2022) to quantify shear stress at different locations on a surface relative to the point of sanitizer spray impingement. Validation was done with experimental flow pattern and velocity field. The geometry was plotted based on the exact components and dimensions of the bench-scale experimental setup as described in *section 2.1* to simulate the sanitizer spray and flow from the nozzle tip to the stainless steel surface. A control-volume approach was employed to integrate the governing equation over each cell in the mesh, and to allow for a set of discrete algebraic equations to be solved for the flow variables, including the pressure, velocity and shear stress.

#### 2.8.1 Governing equations and simplifications in geometry

To avoid complex geometry creation and many mesh grids in simulation, the sanitizer flow was simplified to a high-speed sanitizer column exiting from a custom-shaped pressurized outlet and impinging to the stainless steel wall. Volume of fraction (VOF) module was selected in the simulation since the scenario was simplified as liquid (sanitizer) entering a gas zone (air). For the high-speed injection regarding turbulent flow, a *k-ε* turbulence model was selected for solving the constructed model, with specific equations as follows:

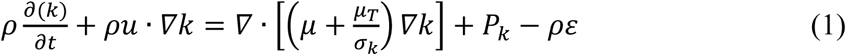

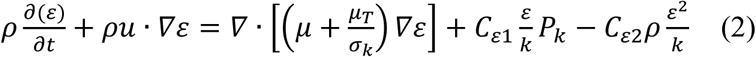

The standard *k-ε* turbulence model was solved using a pressure-based, double-precision implicit solver with second-order discretization for pressure and momentum. The convergence of the simulation was monitored by the continuity residual, residual of the turbulent kinetic energy (*k*), and residual of the turbulent dissipation (*ε*). Convergence was determined based on a calculated residual less than the tolerance of 1 × 10^−4^.

#### 2.8.2 Mesh generation

Fluent generated poly-hexacore mesh was applied to the whole domain. The mesh size was refined until the solved liquid velocity reached a difference below 0.5% for finalization. The maximum and minimum cell size was 0.003 and 0.002 m, respectively. At the pressure inlet, the mesh was refined to a maximum size of 0.002 m and a minimum size of 0.00025 m for converging. A 4-layer boundary layer was set at the wall with the excess ratio set to 0.272 and the growth rate set to 1.2. The computed dimensionless wall distance (*y^+^*) was less than 5 and confirmed that the designed mesh size was appropriate to apply the standard *k-ε* turbulence model for solving the wall shear stress^22^. The total number of cells generated in the whole domain was estimated as 2,881,612.

#### 2.8.3 Initial and boundary conditions

The flow boundary condition at the outlet of the nozzle was set as “pressure inlet” with a gauge pressure of 50 psi (3.45×10^5^ Pa) according to the experimental setup. The nozzle surface and stainless steel coupon surface were set as “wall,” and all other boundaries were set as “pressure outlet” with a gauge pressure of *P* = 0. Non-slip boundary condition was applied, and the properties of the sanitizer solution were set as the built-in definition of liquid water in the software since the dilute solutions have similar flow properties as water at room temperature (Table 1)^23^. In the VOF model, the initial content of the spraying chamber was set as “gas” phase (*v* = 1), and the liquid column within the nozzle was set as “liquid” phase (*v* = 0).

**Table 1.** Fluid properties of air and liquid sanitizer in CFD modeling (25°C)

| Parameters | Species | Value |
| --- | --- | --- |
| Viscosity | Air | $10^{-6}$ Pa·s |
| | Sanitizer (100 ppm) | $10^{-3}$ Pa·s |
| Density | Air | 1.003 kg/m<sup>3</sup> |
|  | Sanitizer (100 ppm) | 998 kg/m<sup>3</sup> |
| Contact angle | Between air and liquid sanitizer | 0.072 N·m |

#### 2.8.4 Post-processing

After convergence, the velocity of sanitizer liquid distribution and the shear stress on the coupon were analyzed directly from ANSYS. Velocity profiles (*u*) of sanitizer were obtained from the top and side view over the chamber as a 2-D contour plot, and the shear stress (*τ_w_*) over *x-* and *y-*direction on the coupon surface were obtained and explored as a 1-D line plot.

#### 2.8.5 Model validation

A high-speed camera (FASTCAM, Photron, San Diego, USA) was utilized to capture the velocity of sanitizer flow during treatment, and data was used to verify the effectiveness of the CFD model. The top view of the flow was captured consecutively, and the image was analyzed with “ImageJ” software (NIH image, USA) for estimating the flow velocity and to compare to the simulation results. The spray angle and spray width on the coupon surface were also obtained from both experiment and simulation, and alignment between those results also supported model validation.

### 2.9 Statistical analysis

All statistical analyses were conducted in RStudio (R version 4.4.3). A one-way ANOVA, followed by Tukey’s HSD test, was used to compare the log reductions achieved by the different researchers during the pilot-scale sanitizing process. One-way Welch’s ANOVA, followed by Games-Howell post-hoc analysis, was performed to compare log reductions between the “Impingement,” “Impingement-adjacent,” and “Fluid-film” locations, as well as to assess the effects of different application and contact time parameters. Welch’s ANOVA and Games-Howell post-hoc analysis were also used to compare static sanitizer application with the spray application methodologies. Welch’s ANOVA and Games-Howell post-hoc analysis were chosen for these comparisons due to significant differences in the magnitude of variances among the groups of interest which violated the homogeneity of variance assumption required for standard one-way ANOVA and Tukey’s HSD test^24^.

## 3. RESULTS AND DISCUSSION

### 3.1 Untrained sanitation operators varied in their application technique and achieved an average Listeria reduction of 2.6 log CFU

Sanitizer efficacy was evaluated under pilot-scale conditions with three untrained researchers to capture variability in application methods and resulting microbial reductions. Spray pattern, application time, and application distance all varied among the researchers (Figure 1B), as did the level of microbial reduction achieved (p < 0.05) (Table 2). Researcher 1 had the shortest application time (6 s), applied sanitizer from the furthest nozzle distance (150 cm), and achieved the lowest microbial reduction (1.9 ± 0.5 log CFU/surface). Researcher 2 applied sanitizer for the longest duration (15 s) with the shortest application distance (45 cm) and achieved the largest microbial reduction of 3.7 ± 0.7 log CFU/surface. Researcher 3 applied sanitizer for 13 s with an application distance of 75 cm, resulting in a reduction of 2.1 ± 0.6 log CFU/surface (Table 2). One-way ANOVA followed by a subsequent Tukey’s HSD test indicated significant differences in microbial reduction among all three of the researchers (p < 0.05) (Table 2).

**Figure 1.**
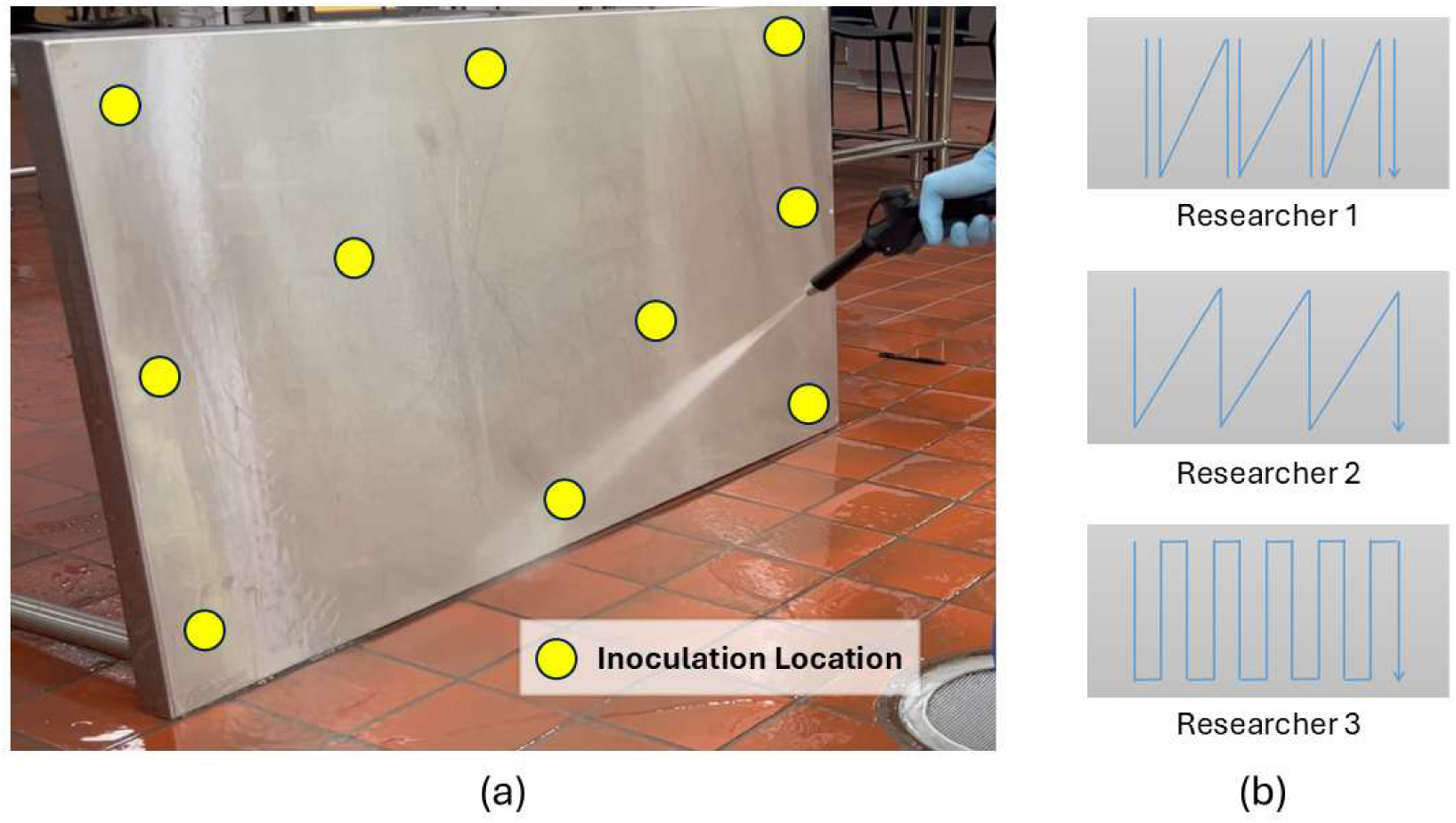
(a) Experimental set-up for pilot-scale sanitization experiment, denoting the inoculation locations (n=10) displayed in yellow. (b) Spray paths for Researcher 1, Researcher 2, and Researcher 3.

**Table 2.** Treatment parameters and outcomes for 3 researchers operating the sprayer for sanitizing. Values in a column denoted with different letters are statistically different from one another (p < 0.05; ANOVA; Tukey’s HSD)

| Researcher | Spray Time<br>(s) | Contact<br>Time (s) | Nozzle<br>Distance<br>(cm) | Average<br>log Reduction (SE) |
| --- | --- | --- | --- | --- |
| 1 | 6 | 120 | 150 | 1.9 (0.5)<sup>A</sup> |
| 2 | 15 | 120 | 45 | 3.7 (0.7)<sup>B</sup> |
| 3 | 13 | 120 | 75 | 2.1 (0.6)<sup>C</sup> |

*3.2 Shear stress was highest near the point of impingement during simulated sanitizer spraying of a flat surface*

The average microbial reduction achieved by all researchers was well below the 5-log standard for EPA registered sanitizers despite an ideal test surface without niches and complete surface coverage with sanitizer being achieved by all spray patterns employed by the researchers^12^. These results indicate that meaningful shortcomings and variability can occur during relatively simple sanitation operations. Indeed, past studies enumerating bacterial populations in facilities before and after sanitation operations often find minimal microbial reduction especially in difficult to clean areas^25,26^. Failure to achieve 5-log reductions can consequently result in food manufacturing facilities failing to meet surface hygiene criteria, such as ≤100 CFU/25 cm² of mesophilic aerobic bacteria ^27–29^.

The significant differences in microbial reduction achieved by the researchers highlights the impacts that application time, spray distance, and spray pattern can have on sanitation efficacy. Past work has shown that increased contact time with sanitizer can enhance microbial inactivation on food contact surfaces^30,31^. Additionally, Scott et al. (1981) found that decreasing application distance significantly improved cleaning efficacy of stainless steel^32^. However, in the current work, even the operator applying sanitizer for the longest duration (15 s), from the shortest application distance (45 cm) did not yield a 5-log reduction on the test surface. Longer application time may improve microbial reduction through enhanced chemical inactivation and physical removal of cells. On the other hand, changes in distance between the sprayer and the surface, as well as the choice of spray pattern, likely primarily affect physical removal of cells through changes in the shear force applied to the surface from the sanitizer spray and flow.

### 3.2 Shear stress was highest near the point of impingement during simulated sanitizer spraying of a flat surface

Shear stress estimation at the “Impingement,” “Impingement-adjacent,” and “Fluid-film” sampling locations via CFD are shown in Figure 2B and Figure 2C. Shear stress was greatest at the “Impingement-adjacent” position at 75 Pa, followed by the “Impingement” location at 25 Pa, and lastly the “Fluid-film” position at 4 Pa. Although the force of the spray was greatest at the point of impingement, shear stress was greater directly adjacent to the impingement point (Figure 2B&C). This is typical of shear stress measurements around the point of impingement in similar systems due to the reduced velocity of the gas or liquid directly at the point of the impingement on the surface^33^. The shear stress being lowest at the “Fluid-film” location was anticipated, as fluid velocity at this location was lower than at the “Impingement” and “Impingement-adjacent” sampling positions. Prior studies confirm that fluid-films on surfaces from falling impinging streams generate less wall shear stress^9^. The low shear stress on surfaces that do not get directly sprayed may have implications for the efficacy of sanitation operations. As previously shown, the spray patterns of operators were highly variable and resulted in uneven application of the sanitizer stream and impingement point on a test surface (Figure 1B).

**Figure 2.**
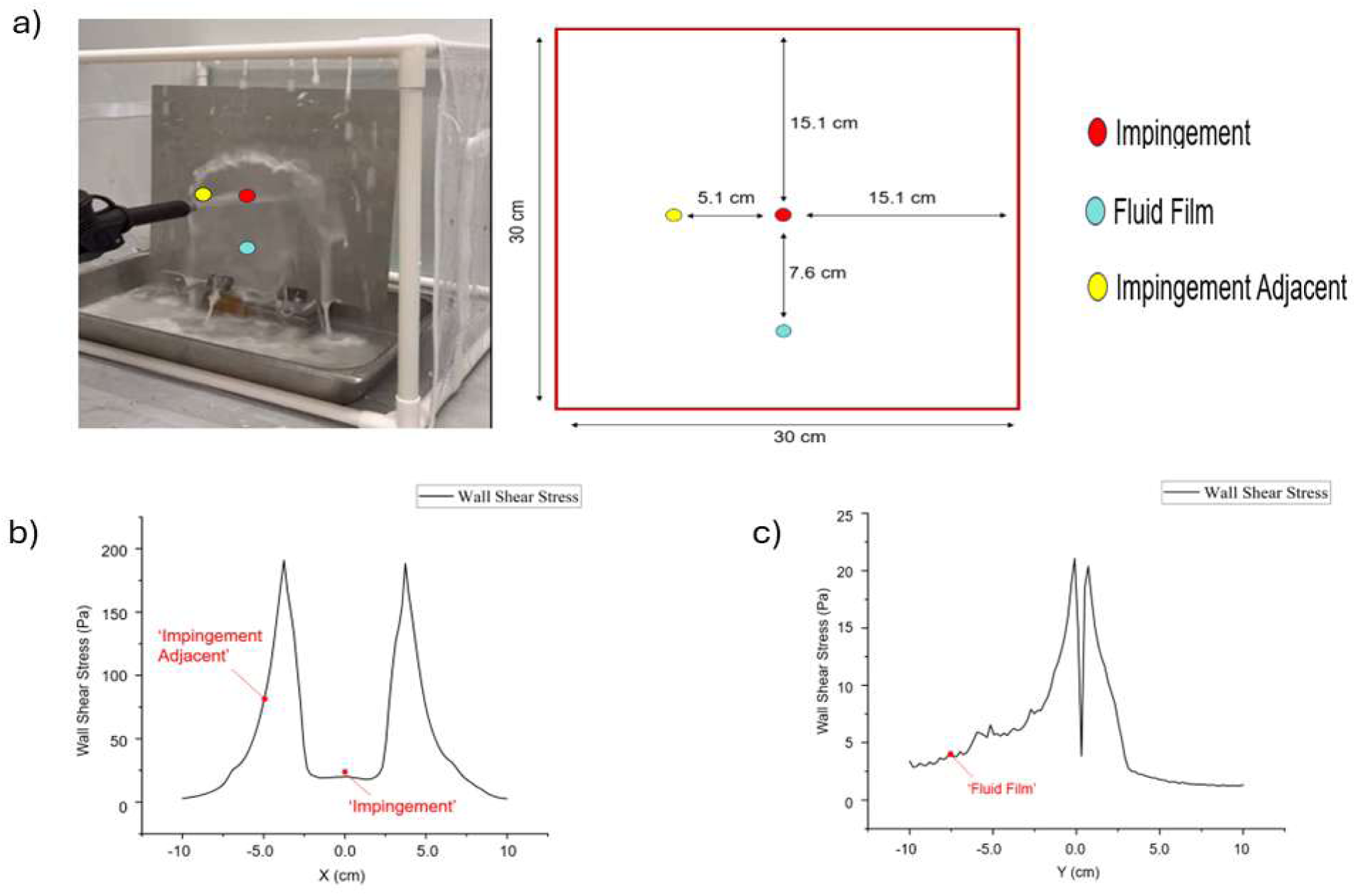
(a) Bench-scale spray-chamber configuration with inoculation sites. CFD simulation results for the (b) top view of the shear stress profile and (c) side view of shear stress profile.

### 3.3 Microbial reduction from spraying was greatest at, and adjacent to, the point of impingement for both water and sanitizer sprays

During rapid exposure (3 s application time – 0 s extended contact time after cessation of spraying), average microbial reduction via a commercial sprayer in the bench-scale spray-chamber was the greatest at the “Impingement” sampling location followed by the “Impingement-adjacent” location and lowest at the “Fluid-film” position for both the water and sanitizer treatments (Table 3). The log reduction from spray application of water at the “Impingement” position was 3.4 ± 0.6 log CFU/surface, at the “Impingement-adjacent” position was 3.2 ± 0.5 log CFU/surface, and at the “Fluid-film” position was 0.4 ± 0.8 log CFU/surface.

**Table 3.** log reductions of *Listeria innocua* from bench-scale spray-chamber trials (3 s application time, 0 s contact time) and associated shear stress values from CFD simulations. Values in a column not connected by the same letter are statistically different from one another (p < 0.05; Welch’s-ANOVA; Games-Howell)

| Site | Treatment | Shear Stress (Pa) | log Reduction (SE) |
| --- | --- | --- | --- |
| Impingement | Sanitizer | 25 | 7.5 (0.5)<sup>C</sup> |
| Impingement-adjacent | Sanitizer | 75 | 6.4 (0.7)<sup>AC</sup> |
| Fluid-film | Sanitizer | 4 | 0.4 (0.1)<sup>D</sup> |
| Impingement | Water | 25 | 3.4 (0.6)<sup>AB</sup> |
| Impingement-adjacent | Water | 75 | 3.2 (0.5)<sup>ABD</sup> |
| Fluid-film | Water | 4 | 0.4 (0.8)<sup>BD</sup> |

Although the mean reductions between these treatments was > 3 log CFU in some instances, differences between locations following water spray were not statistically significant (p > 0.05; Welch’s ANOVA; Games-Howell). This lack of statistically significant differences between treatments despite the large difference in means may be attributable to high intra-treatment variability as well as the multiple comparisons adjustment performed to avoid inflating type-I error.

For surfaces treated by the sprayer with 100 ppm of pressurized sanitizer, the log reduction was again highest at the “Impingement” location followed by the “Impingement-adjacent” and “Fluid-film” locations at 7.5 ± 0.5 log CFU/surface, 6.4 ± 0.7 log CFU/surface, and 0.4±0.1 log CFU/surface, respectively (Table 3). Welch’s ANOVA followed by Games-Howell post-hoc analysis indicated a significant difference in microbial reduction from the sanitizer treatment between the “Fluid-film” position and the other two sampling locations (p < 0.05) (Table 3). Although the microbial reductions at the “Impingement” and “Impingement-adjacent” sampling locations were greater with sanitizer than with water, the >3-log reductions at these locations achieved by water alone demonstrate a significant portion of the reduction achieved was attributable to physical removal rather than chemical inactivation.

The “Fluid-film” location had the lowest microbial reduction during spray application of both water and sanitizer, likely due to the minimal shear forces at this location. This low reduction is unlikely to be attributable to deposition of microorganisms detached from the “Impingement” or “Impingement-adjacent” positions, as any cells in the flowing sanitizer would be inactivated while suspended in solution^34^. In contrast, it was unexpected that microbial reduction was highest (although not significantly different) at the “Impingement” location despite higher modeled shear stresses at the “Impingement-adjacent” location. This could be due to larger normal forces from the sanitizer spray directly at the point of impingement which may result in removal of more cells. It is also possible that the simulation model, which simulated shear stress under ideal flow conditions, underestimated the shear stress at the site of impingement due to deviations from CFD predicted flow that occur in the physical world^35^. Alternatively, it may be the case that microbial reduction peaks after a certain level of shear force, which would explain why there was not a significant difference in microbial reduction between the “Impingement” and “Impingement-adjacent” locations. Indeed, Gibson et al. (1999) reported approximately a 3 log CFU/cm^2^ and 2 log CFU/cm^2^ reduction of surface-adhered *Staphylococcus aureus* and *Pseudomonas aeruginosa* biofilms, respectively, from a stainless steel surface with a pressurized water sprayer system and did not see this value increase after increasing the pressure of the sprayer from 17 to 70 bar^36^.

The large difference in log reductions between surfaces that were in proximity to the point of impingement during sanitizer spraying (7.5 ± 0.5 log CFU/surface at the "Impingement" sampling location) versus those that were not (0.4 ± 0.1 log CFU/surface at the “Fluid-film” sampling location) may partially explain why manual spraying by operators failed to achieve an average 5-log reduction despite having a longer sanitizer contact time (120 s vs 0 s). The highest average manual reduction (3.7 ± 0.7 log CFU/surface) was still well below reductions at both “Impingement” (7.5 ± 0.5 log CFU/surface) and “Impingement-adjacent” (6.4 ± 0.7 log CFU/surface) locations from the bench-scale trial, yet this was still greater than the reduction achieved at the “Fluid-film” location (0.4 ± 0.1 log CFU/surface). Therefore, variability in spray patterns among operators likely resulted in portions of the surface not receiving direct sanitizer impingement and, consequently, experiencing minimal shear stress, which may partially explain the low and variable microbial reductions observed despite similar application times across operators. This provides additional evidence that differences in fluid mechanics as a consequence of manual application can have major impacts on sanitization efficacy.

### 3.4 Microbial reduction increased with longer sanitizer application and contact times, but sanitizer efficacy differed between application methods

The microbial reduction at the “Fluid-film” location increased from 0.4±0.1 log CFU/surface to 2.0 ± 0.1 log CFU/surface when a 120 s contact time with the sanitizer was added following the cessation of the 3 s spray application time (Table 4). Increasing the spray application time from 3 s to 120 s further increased the log reduction per surface at the “Fluid-film” location to 6.3±0.3 log CFU/surface despite a 0 s contact time (p < 0.05) (Table 4). In comparison, submersion of a coupon in static sanitizer for 3 s followed by immediate swabbing and no extra contact time yielded a reduction of 2.3 ±0.1 log CFU/surface, a significantly greater reduction than at the “Fluid-film” sampling location under the same conditions (p < 0.05). Adding a 120 s contact time following coupon submersion for 3 s increased the reduction to 4.4 ± 0.6 log CFU/surface. Submerging the coupon for 120 s and no additional contact time resulted in a reduction of 6.0 ±0.1 log CFU/surface (Table 4).

**Table 4.** *Listeria innocua* reduction on submerged coupons and at the “Fluid-film” location under varying sanitizer application and contact times. Values in a column denoted with different letters are statistically different from one another (p < 0.05; Welch’s-ANOVA; Games-Howell)

| Site | Application Time (s) | Contact Time (s) | log Reduction (SE) |
| --- | --- | --- | --- |
| Submerged coupon | 3 | 0 | $2.3 (0.1)^A$ |
| | 3 | 120 | $4.4 (0.6)^{ABC}$ |
| | 120 | 0 | $6.0 (< 0.1)^{CD}$ |
| Fluid-film | 3 | 0 | $0.4 (0.1)^B$ |
| | 3 | 120 | $2.0 (0.1)^{AC}$ |
| | 120 | 0 | $6.3 (0.3)^D$ |

These results highlight how typical validation of sanitizer treatments often do not reflect the dynamic environments where sanitation occurs, such as in pilot plants or industrial facilities where there may be large variability in the application time and shear stress of sanitizer sprays. For example, coupon submersion for 3 s outperformed manual spraying by operators despite the latter having greater shear forces and longer application time albeit over a larger surface area. This likely occurred because submersion ensured complete and uniform contact of the sanitizer with the entire surface over the application time whereas in the fluid film, uneven flow of sanitizer down the coupon could have resulted in inconsistent coverage of the inoculation site with sanitizer.

Coupon submersion and spraying at the “Impingement” location also overestimated how much microbial reduction was achieved at “Fluid-film” spray locations which can be assumed to accurately reflect the conditions that a meaningful proportion of environmental surfaces achieve during sanitation operations. Baker et al. (2025) similarly found that microbial reductions from application of a superheated steam sanitation treatment on bench-scale surfaces were much higher than what was achieved during pilot-scale application^37^. Training interventions could increase microbial reduction from operator spray application of sanitizer^38^. Emphasis on ensuring surfaces achieve sufficient shear forces to dislodge cells could improve sanitation outcomes. To counteract the lack of shear stress in fluid-films, it may be necessary to increase the time these locations are treated with sanitizer to yield sufficient microbial reductions. Notably, sanitizer exposure at the “Fluid-film” location during spray application, as well as submersion in sanitizer, each achieved a 5-log reduction on stainless steel surfaces when sufficient application time was provided (Figure 3). These improvements may result in microbial reduction values closer to those observed in the bench-scale spray and submerged coupon trials; however, research into training interventions to reduce post-pasteurization product recontamination, an indication of poor sanitation, has shown major limitations^39^.

**Figure 3.**
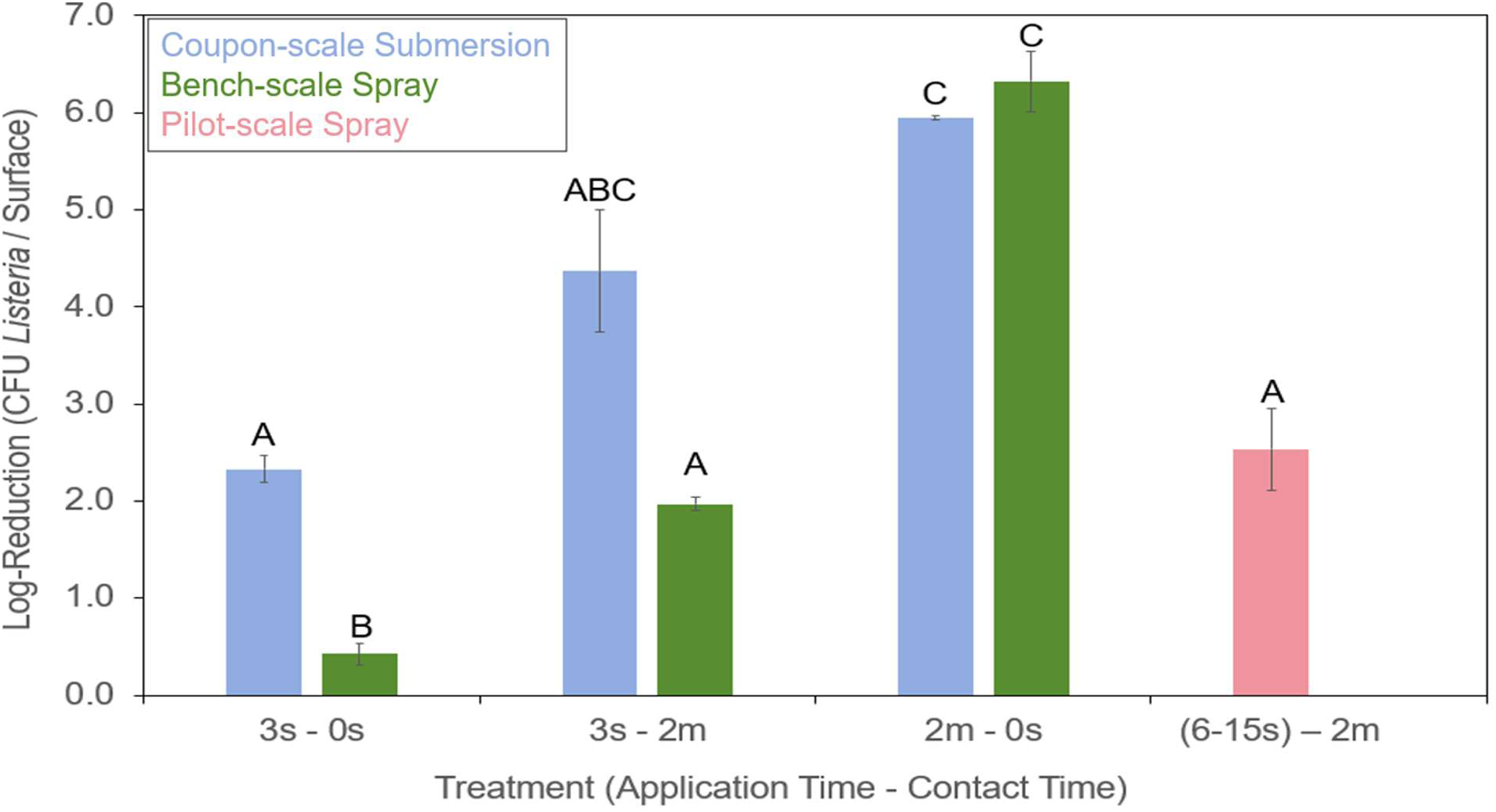
Reduction of *L. innocua* from sanitizer treatment under three conditions: coupon submersion (blue), sprayers in a bench-scale experiment (green), and researchers using sprayers at the pilot-scale (pink). Data from the “Fluid-film” location in the bench-scale experiments is presented in this figure (green). Bars denoted with different letters are statistically different from one another (p < 0.05; Welch’s-ANOVA; Games-Howell)

## 4. CONCLUSION

Our study demonstrates how variability in the physical parameters of sanitizer spraying can result in lower than expected microbial reductions during sanitation. Operators did not achieve a 5-log reduction of *Listeria* even on an easy-to-clean flat stainless steel test surface without niches. This can be partially explained by insufficient application time of sanitizer and highly variable shear stress applied to the surface depending on the spray pattern they chose. Controlled bench-scale experiments revealed that *Listeria* reduction was greatest on surfaces experiencing high shear stress at the "Impingement" and “Impingement-adjacent” sampling locations for both sanitizer and water spray treatments. This emphasizes the contribution of physical cell dislodgment during sanitizing, which is an often overlooked element of sanitation. Microbial reduction was minimal on surfaces only exposed to the “Fluid-film” of sanitizer where shear stress was lowest. However, further experiments showed that even under these conditions a sufficiently long application time of sanitizer (120 s) could achieve a > 5-log reduction/surface. Therefore, experiments measuring the efficacy of sanitizers may overestimate or underestimate sanitation outcomes compared to real world conditions if shear stress and application time of sanitizer are not accounted for. An emphasis on maximizing the shear forces applied on surfaces through the impingement point of sanitizer spray could be a helpful principle for the training of sanitation operators or the development of spraying tools. Additionally, increasing the time spent spraying a surface and allowing for a longer sanitizer contact time after spraying are critical for maximizing microbial reduction. In reality, sanitation outcomes from sanitizer spray application are dynamic and are likely to fall short of achieving >5-log reduction/surface even on flat surfaces let alone within environmental niches. This complicates decision making in operations like low moisture food processing where the risk of introducing water to the environment during sanitization must be balanced against the efficacy of wet sanitation compared to dry sanitation. This work underscores the need for future research to better understand gaps between experimental sanitation validation and real world efficacy of sanitation operations.

